## Supplemental Figure legends for "A modular CRISPR screen identifies individual and combination pathways contributing to HIV-1 latency"

**Supplemental information:**

**Supplemental Table 1**. GuideRNA sequences for sgRNAs of the HuEpi library.

**Supplemental Table 2**. MAGeCK analysis output by sgRNA for the Latency HIV-CRISPR screen.

**Supplemental Table 3**. MAGeCK analysis output by gene or NTC for the Latency HIV-CRISPR screen.

**Supplemental Table 4**. MAGeCK analysis output by sgRNA for the AZD5582 LRA Latency HIV-CRISPR screen.

**Supplemental Table 5**. MAGeCK analysis output by gene or NTC for the AZD5582 LRA Latency HIV-CRISPR screen.

**Supplementary Table 6**. Primers sequences, sgRNA sequences, and next-generation sequencing barcodes.

**Supplemental Figure 1. Validation of the top hit from the Latency HIV-CRISPR screen in primary CD4+ T cells.**
**(A)** Representative flow cytometry plots of viral reactivation levels in wildtype primary CD4+ T cells and primary CD4+ T cell model of HIV latency cells upon knockout of AAVS1 and CUL3.

**Supplemental Figure 2. AZD5582 dose curve in J-Lat cells.**
**(A)** AZD5582 dose curve performed on both J-Lat 10.6 and 5A8 cell lines to determine viral reactivation levels.

**Supplemental Figure 3. The genome-wide signal of automated CUT&Tag replicates is highly correlated.**
(**A**) Correlation Matrix colored according to the pair-wise Pearson correlation of pan-H4Ac, BRD4, and IgG negative control samples across the merged pan-H4Ac and BRD4 peak sets. All pan-H4Ac samples group together by hierarchical clustering as do all of the BRD4 samples. **(B)** Same as (A) but showing the pair-wise Pearson correlation of RNA-Pol2-S5p and RNA-Pol2-S2p over the merged peak sets of these marks.
