## Supplementary figures and images for "A modular CRISPR screen identifies individual and combination pathways contributing to HIV-1 latency"

### Supplemental Figures

#### Supplemental Figure 1

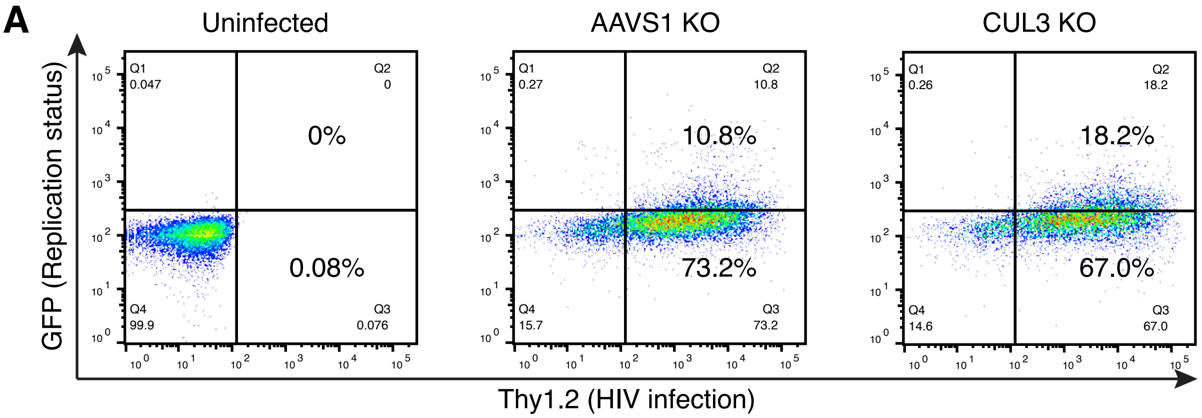

#### Supplemental Figure 2

A

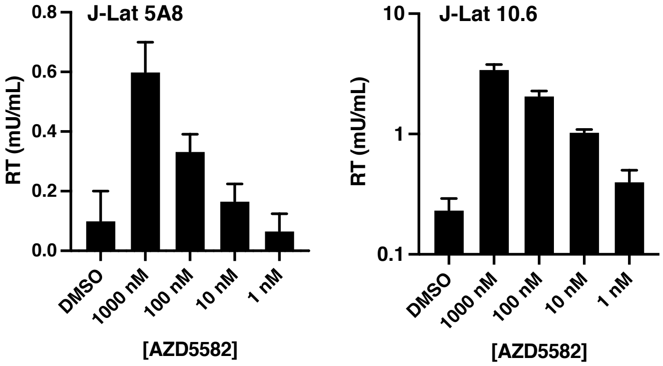

#### Supplemental Figure 3

A

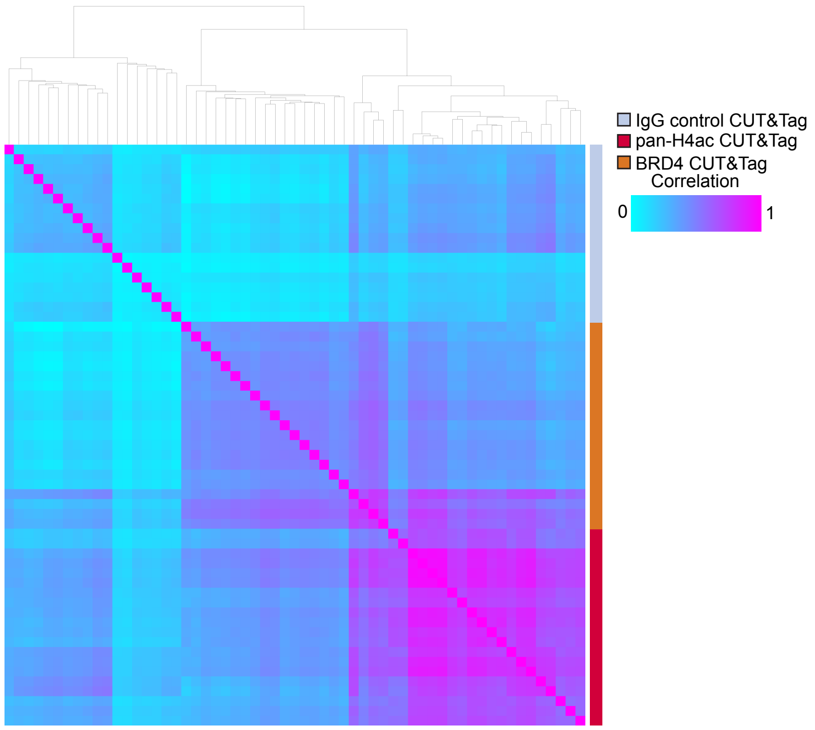


B

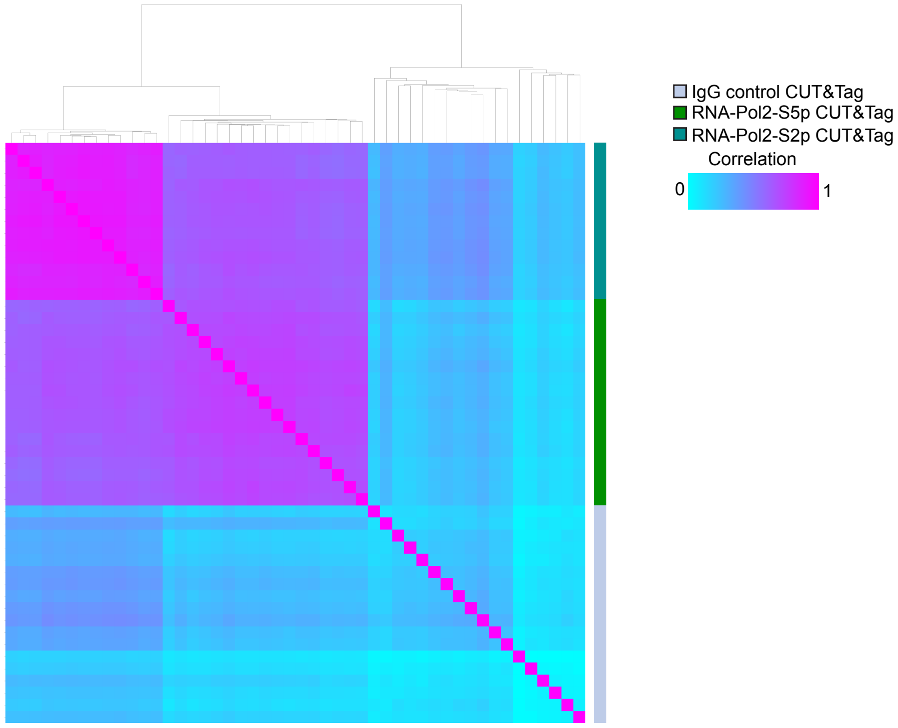
